# NATO3 protects dopaminergic neurons in mouse *in vivo* and human *in vitro* Parkinson’s disease models

**DOI:** 10.1101/2025.10.10.681573

**Authors:** Eva P. Valencia-Alarćon, Lou C. Duret, Emi Kobayashi, Olivia Cattaneo, Bernard L. Schneider, Emi Nagoshi

## Abstract

Parkinson’s disease (PD) is a devastating neurodegenerative disorder primarily characterized by the progressive and unstoppable loss of dopaminergic (DA) neurons in the substantia nigra. We previously identified NATO3 (FERD3L), a conserved developmental transcription factor, as essential for maintaining DA neuron function during aging. Here, we show that AAV-mediated *Nato3* gene transfer into the mouse substantia nigra prevents DA neuron degeneration in both MPTP-induced and α-synuclein (α-Syn) overexpression PD models. This neuroprotective effect is achieved by improving autophagic flux and *⍺*-Syn clearance. Furthermore, lentiviral-mediated NATO3 overexpression in human midbrain DA neurons, derived from induced pluripotent stem cells carrying the pathological *⍺*-Syn A53T mutation, effectively reversed key disease hallmarks. These include *⍺*-Syn accumulation, aberrant mitochondrial morphology, autophagic impairments, and compromised neurite structure. Collectively, these *in vivo* and *in vitro* findings highlight NATO3’s role in safeguarding DA neurons against pathological cellular events, positioning NATO3 as a therapeutic target for PD.

## Introduction

Parkinson’s disease (PD) is the second most common neurodegenerative disorder and the leading cause of motor dysfunction. Its hallmark motor symptoms are largely attributed to the progressive loss of dopaminergic (DA) neurons in the substantia nigra pars compacta, resulting in significant DA deficits in the striatum. The age-dependent progression of this neuronal loss is currently irreversible, posing a major challenge to an aging global population.

The pathogenesis of both sporadic and familial PD is complex and multifactorial, involving numerous genetic risk loci and pathogenic mutations. A unifying feature observed in both types of the disease is the accumulation of misfolded proteins, most notably α-synuclein (α-Syn) – the primary component of the cytoplasmic inclusions known as Lewy bodies. Preclinical studies using different models suggest that enhancing protein homeostasis and the clearance of protein aggregates may be a promising therapeutic strategy (Calabresi *et al*., 2023; Sharma *et al*., 2025). Similarly, impaired mitochondrial function is a central pathological event in PD, highlighting the importance of efficient mitophagy as a neuroprotective mechanism (Ge *et al*., 2020; Narendra & Youle, 2024). However, the interconnected nature of these pathways, with numerous molecular players, presents a significant challenge. Targeting a single player may not only fail to improve the clearance of aberrant protein species but could also have confounding effects (Khan *et al*., 2024; Luo, 2025).

Several transcription factors regulating the development of midbrain DA neurons, such as NURR1, FOXA1/2, EN1/2, PITX3, OTX2, and LMX1A/B, continue to be expressed in adult DA neurons. These factors play crucial roles in maintaining these neurons in adult animals (Blaudin de The et al., 2016; Di Salvio et al., 2010; Domanskyi et al., 2014; Doucet-Beaupre et al., 2016; Kadkhodaei et al., 2009; Kittappa et al., 2007; Ariadna Laguna et al., 2015; van den Munckhof et al., 2003). Despite their importance, the mechanisms underlying their neuroprotective functions are not yet fully understood. Likewise, our previous work has established that *Nato3* (*Ferd3l*), encoding a bHLH-transcription factor, falls into this category of developmental transcription factors crucial for maintaining DA neuron function in adulthood (Miozzo *et al*., 2022). In vertebrates, NATO3 is expressed in developing mesencephalic floor plate cells (the progenitors of midbrain DA neurons) and promotes neurogenesis of mesencephalic dopaminergic neurons through suppressing *Hes1* (Ono *et al*., 2010). While the SHH-FOXA2 axis activates NATO3 expression, NATO3 also feedback-upregulates these genes (Nissim-Eliraz *et al*., 2013). Additionally, NATO3 has been shown to upregulate the expression of genes that specify DA neuron cell fate, including *Lmx1b* (Peterson *et al*., 2019). We have shown that the conditional ablation of *Nato3* in post-differentiated DA neurons impairs mitochondrial integrity in the midbrain DA neurons in aged mice. Importantly, the role of *Nato3* in maintaining mitochondrial integrity within DA neurons is conserved from invertebrate species, particularly *C. elegans* and *Drosophila* (Bou Dib *et al*., 2014; Miozzo *et al*., 2022). However, the downstream mechanism by which Nato3 maintains mitochondrial quality in DA neurons remains unknown.

Notably, while loss of function in *Fer2*, the *Drosophila Nato3* homolog, increases susceptibility to oxidative stress and induces age-dependent degeneration of DA neurons, its overexpression prevents DA neuron degeneration in genetic- and neurotoxin-induced PD models (Bou Dib *et al*., 2014; Tas *et al*., 2018; Miozzo *et al*., 2022). However, whether the overexpression of NATO3 has similar neuroprotective potential in mammals remains to be elucidated. This work bridges this knowledge gap by investigating the effect of NATO3 in mice and human midbrain DA (mDA) neurons in the context of PD and its underlying mechanisms. We employ AAV-mediated NATO3 overexpression in the SN of two mouse models of PD, MPTP-induced and *⍺*-Syn overexpression. We further use lentiviral-mediated NATO3 overexpression in human mDA neurons differentiated from induced pluripotent stem cells (iPSCs) carrying the pathogenic α-Syn A53T mutation, to evaluate its ability to mitigate PD-related pathological characteristics.

## Results

### *Nato3* gene delivery prevents degeneration of DA neurons in mouse models of PD

To test if *Nato3* overexpression confers dopaminergic neuroprotection, we utilized two well-characterized mouse models of PD: the 1-methyl-4-phenyl-1,2,3,6-tetrahydropyridine (MPTP) model and a model for *⍺*-synuclein (*⍺*-Syn)-linked PD. For *Nato3* overexpression, we generated the AAV6-pgk-*mNato3*-WPRE vector (AAV-Nato3), which is designed to express mouse *Nato3* (*mNato3*) under the control of the phosphoglycerate kinase 1 (pgk1) promoter. The woodchuck hepatitis virus posttranscriptional regulatory element (WPRE) is added before a polyadenylation signal to increase the expression levels of *mNato3* (Higashimoto *et al*., 2007).

MPTP is a mitochondrial toxin that causes the selective loss of DA neurons, recapitulating a major pathophysiology of PD (Meredith & Rademacher, 2011). This makes MPTP administration a widely used method to model PD, although inconsistent effects on behavior have been reported (Santoro *et al*., 2023). To investigate the effect of *Nato3* overexpression on MPTP-induced DA neuron loss, we delivered AAV-Nato3 via stereotactic injection into one side of the SN in 3-month-old mice. One week after the stereotactic injection, the mice received intraperitoneal (IP) injections of MPTP (20mg/Kg) daily for 5 consecutive days. IP injections of saline solution and a stereotactic injection of the vehicle (PBS) were used as controls for MPTP and AAV-Nato3, respectively. The number of DA neurons in the SN was analyzed one month after the MPTP treatment with anti-tyrosine hydroxylase (TH) immunostaining (Fig. 1a). We found that AAV-Nato3 stereotactic injection effectively prevented the MPTP-induced DA neuron loss in the SN (Figs. 1b and c).

**Figure 1.**
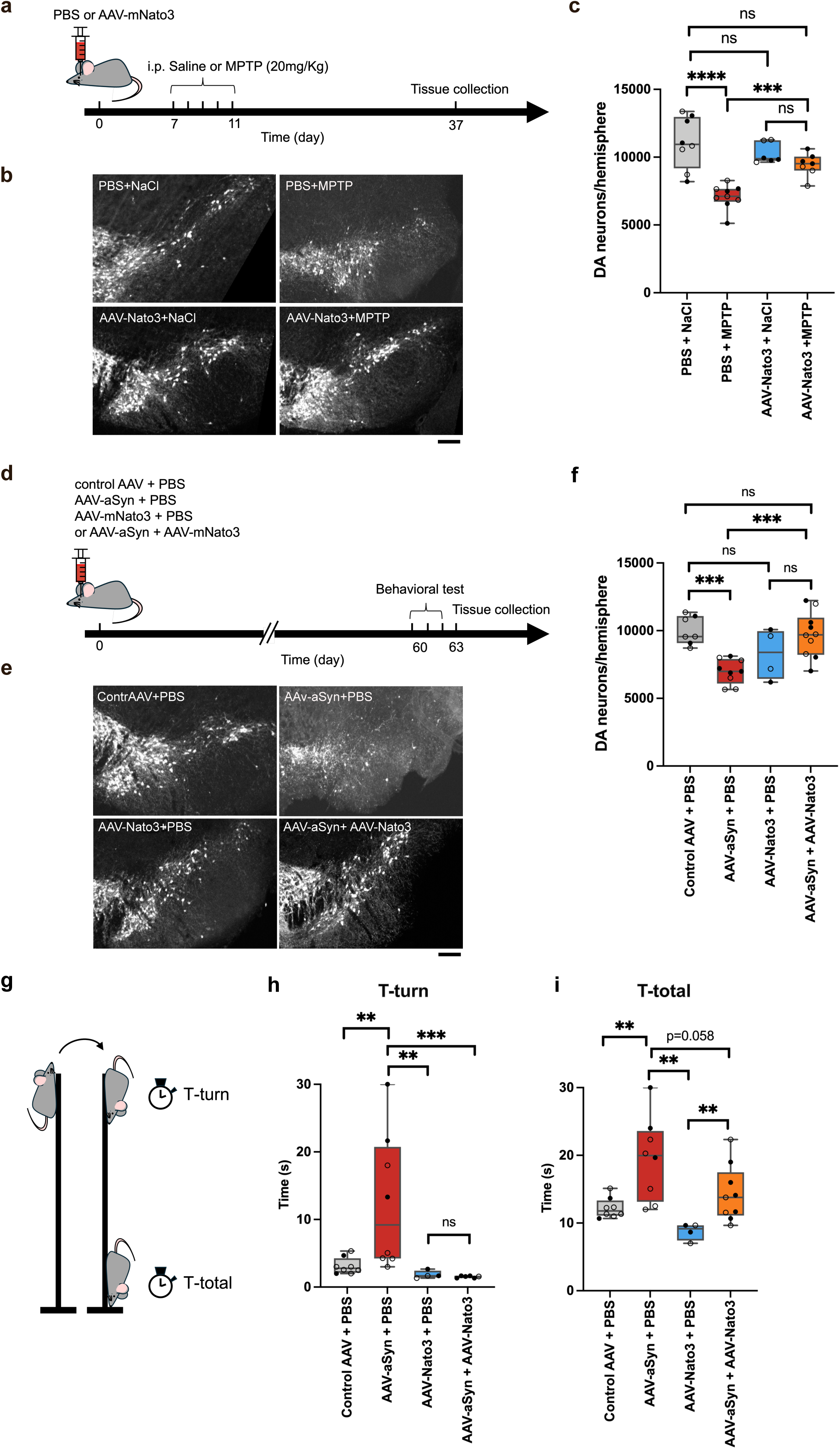
*Nato3* gene delivery protects against degeneration of DA neurons in mouse models of PD. **a.** Schematic of the experimental procedure for testing the effect of *Nato3* gene delivery in MPTP-treated mice. **b**. Representative images of the SN analyzed one month post-injection on the ipsilateral side, with immunostaining for TH. Scale bar, 200 µm. **c.** Quantification of DA neurons in the ipsilateral SN indicates the protective effect of AAV-Nato3 against MPTP-induced loss of DA neurons. n=6–9 per group. **d.** Schematic of the experimental timeline to test the effect of *Nato3* gene delivery on neuropathology induced by AAV7-mediated α-Syn overexpression. **e.** Representative images of the SN, analyzed two months post-injection on the ipsilateral hemisphere, immunostained for TH. Scale bar, 200 µm. **f.** Quantification of DA neurons in the ipsilateral SN indicates the protective effect of AAV6-Nato3 against degeneration of DA neurons induced by α-Syn overexpression. n=4–10 per group. **g-i.** The pole test measures the time taken to orient downward (T-turn) and the total time to turn and descend the pole (T-total). *Nato3* gene delivery prevents the increase in T-turn induced by α-Syn overexpression (**h**) and shows a tendency to ameliorate the increase in T-total (**i**). n=4–10 per group. In **c, f, h,** and **i,** the center line of the box plot represents the median, the box boundaries are the 25th and 75th percentiles, and the whiskers indicate the minimum and maximum values. The closed circles represent values from males, and the open circles represent females. Statistical comparisons were performed using a two-tailed Mann-Whitney U test. **p<0.01, ***p<0.001, and ****p<0.0001. ns, not significant.

Previous research has established that AAV vector-mediated overexpression of α-Syn in the SN causes a progressive loss of DA neurons in the injected site, accompanied by the accumulation of α-Syn-positive aggregates and locomotor impairments (Oliveras-Salvá *et al*., 2013). Following this prior work, we generated the AAV vector expressing α-Syn under the control of the neuron-specific *synapsin 1* (syn1) promoter (AAV7-syn1-*⍺*Syn WT-WPRE; AAV-*⍺*Syn) and a control vector that does not express any gene (AAV7-syn1-empty-WPRE). To determine the effect of *Nato3* overexpression in this model, AAV-*⍺*Syn was stereotactically co-injected into one side of the SN with its PBS vehicle or with AAV-Nato3 in 3-month-old mice. Control animals were injected with the control vector in PBS or AAV-Nato3 in PBS. Two months post-injection, locomotor behavior was analyzed using a pole test, an assay suitable for evaluating locomotor impairments associated with reduced striatal dopamine (Matsuura *et al*., 1997). Subsequently, the viability of DA neurons in the substantia nigra was examined with anti-tyrosine TH immunohistochemistry (Fig. 1d).

As expected, AAV-*⍺*Syn stereotactic injection led to a significant nigral DA neuron loss in the SN of the injected hemisphere (Supplementary Figs. 1a-c, Figs. 1e and f). In the pole test, these mice exhibited significantly longer times to turn (T-turn) and descend the pole (T-total), compared to mice injected with the control AAV (Figs. 1g-i). These behavioral results confirm that AAV-mediated *⍺*-Syn overexpression also induces locomotor impairments. Crucially, AAV-Nato3 co-injection with AAV-*⍺*Syn prevented this *⍺*Syn-induced DA neuron loss (Figs. 1e and f). The co-injection also significantly attenuated the *⍺*Syn-induced locomotor impairments, specifically the increased T-turn duration (Fig. 1h). The T-total duration also showed a trend towards improvement with the co-injection of AAV-Nato3, although statistical significance was not reached. This may be partly attributed to the high variability in the pole test performance among the AAV-*⍺*Syn-injected mice.

### *Nato3* promotes the autophagy-lysosomal pathway and the clearance of α-Syn aggregates

Given that *Drosophila Fer2* is essential for maintaining autophagic flux in midbrain DA neurons (Tas *et al*., 2018), we asked if *Nato3* is similarly crucial for the autophagy-lysosomal pathway in mammals. We therefore investigated whether the autophagic flux is impaired in mice with conditional ablation of *Nato3* in differentiated DA neurons (*Nato3* cKO mice) (Miozzo *et al*., 2022). Immunohistochemical analysis of P62/SQSTM1, a selective autophagy receptor that delivers substrates into autophagic vesicles (Fig. 2a) (Vargas *et al*., 2023), revealed elevated P62 levels within the SN DA neurons of aged *Nato3* cKO mice compared to controls. This finding indicates that *Nato3* ablation blocks autophagic flux, supporting its essential role in maintaining the autophagy-lysosomal pathway (Figs. 2b and c).

**Figure 2.**
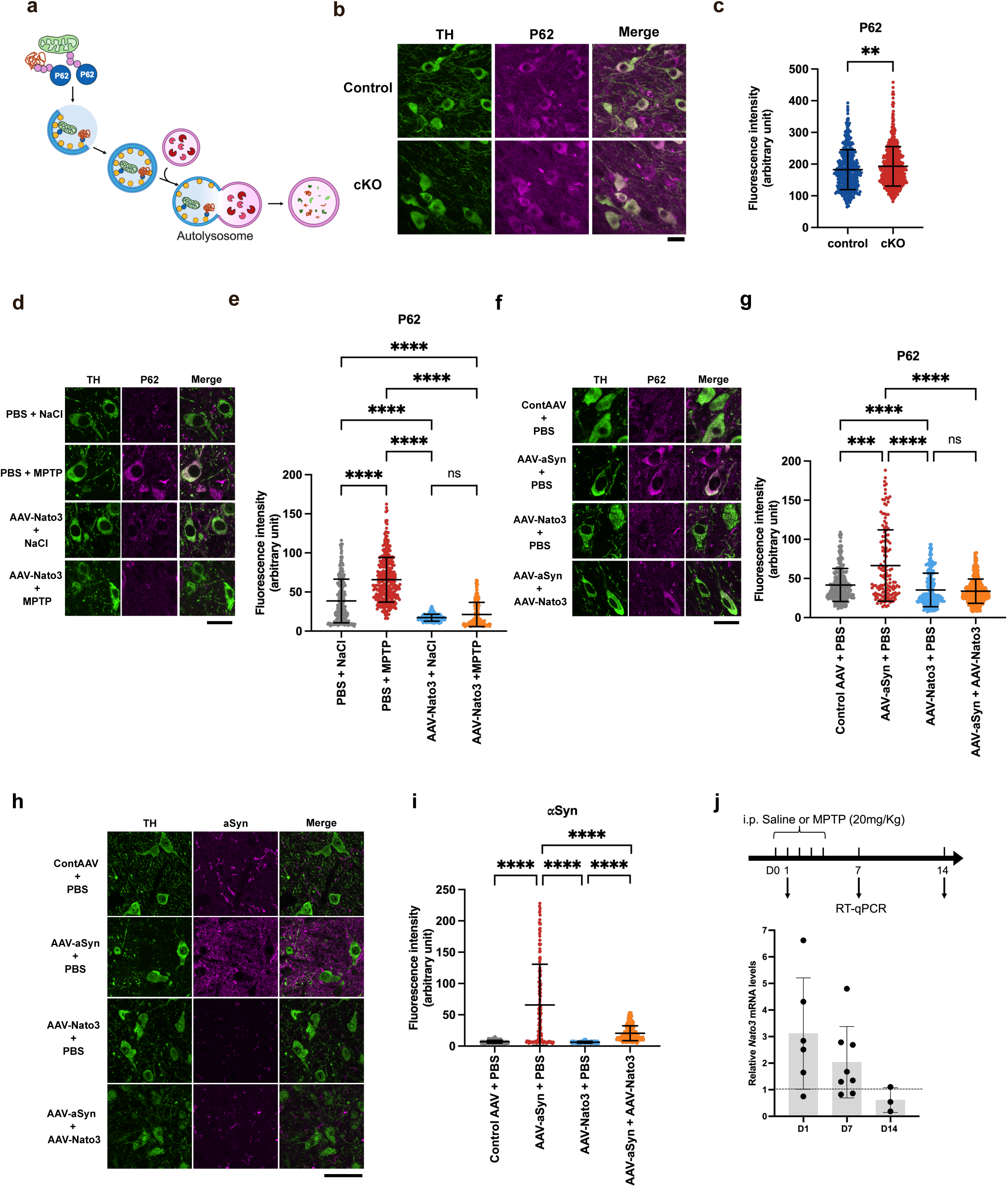
*Nato3* stimulates autophagic flux and the clearance of accumulated α-Syn. **a.** Schematic of the autophagy-lysosomal pathway. P62/ SQSTM1 protein is a selective autophagy receptor. **b.** Representative confocal images of the SN from 26- to 32-month-old control and *Nato3* cKO mice immunostained for TH and P62. Scale bar, 20 µm. **c.** Quantification of P62 fluorescence intensity in the DA neurons. n=3 mice per group. **d.** Representative confocal images of the SN sections immunostained for TH and P62. Mice received PBS or AAV-Nato3 stereotactic injection, followed by saline or MPTP ip injections. Scale bar, 20 µm. **e.** Quantification of P62 fluorescence intensity in the DA neurons demonstrates that *Nato3* gene delivery mitigates the increase in P62 levels induced by MPTP. n=6–9 mice per group. **f.** Representative confocal images of the SN sections immunostained for TH and P62 from the mice injected with control AAV7 alone or AAV7-αSyn alone, as compared to mice injected with either AAV6-Nato3 alone or AAV6-Nato3 plus AAV7-αSyn. Scale bar, 20 µm. **g.** Quantification of P62 fluorescence intensity in the SN DA neurons, indicating that AAV-Nato3 prevented the increase in P62 levels induced by AAV-αSyn. n=4–10 mice per group. **h.** Representative confocal images of the SN sections immunostained for α-Syn and TH from the mice injected with control AAV7 alone or AAV7-αSyn alone, as compared to mice injected with either AAV6-Nato3 alone or AAV6-Nato3 plus AAV7-αSyn. Scale bar, 20 µm. **i.** Quantification of α-Syn fluorescence intensity in the SN DA neurons. AAV6-Nato3 co-injection mitigated the accumulation of α-Syn induced by AAV7-αSyn. n=4–10 mice per group. **j.** MPTP treatment upregulates *Nato3* expression in the SN. *Nato3* mRNA levels in the SN were quantified by RT-qPCR in mice treated with MPTP and are shown relative to the saline control group (dotted line). Mean ± SD. n=3–8. In **c, e, g,** and **i,** scatter plots show mean ± SD. Each data point represents the value from a TH-positive cell. A two-tailed Mann-Whitney test was used for statistical comparison. **p<0.01, ***p<0.001, and ****p<0.0001. ns, not significant.

Having established that *Nato3* overexpression can confer protection on nigral DA neurons from degeneration in both the MPTP and *⍺*Syn models, we next investigated if the neuroprotective role of *Nato3* is associated with its role in regulating the autophagy-lysosomal pathway. Immunohistochemistry of the brain sections one month post-MPTP injections revealed a significant increase in P62 levels in the SN DA neurons (Figs. 2d and e). This finding indicates the blockage of autophagic flux in the MPTP-treated mice, concordant with the accumulating evidence of autophagic impairments in PD patients (Moors *et al*., 2017; Ma *et al*., 2019). AAV-Nato3 stereotactic injection prior to the MPTP IP injections effectively reduced the P62 levels, indicating the improvement in autophagic flux (Figs 2d and e). It is also noteworthy that AAV-Nato3 with saline injection reduces P62 levels even compared to the control treatment (PBS and saline), suggesting that an increase in Nato3 generally improves autophagic flux (Figs 2d and e).

Similarly, P62 levels were increased in the nigral DA neurons two months post-stereotactic injection of AAV-*⍺*Syn (Figs. 2f and g). These findings are congruent with a previous study indicating the causal role of autophagy flux blockages in Lewy body pathology (Sato *et al*., 2018). Importantly, co-injection of AAV-Nato3 reduced these elevated P62 levels (Figs 2f and g). Furthermore, immunostaining for *⍺*-Syn revealed a significant increase in *⍺*Syn levels following AAV-*⍺*Syn stereotactic injection, which was notably decreased by the co-injection of AAV-Nato3 (Figs. 2h and i). Collectively, these results indicate that *Nato3* gene delivery improves autophagic flux, which in turn mitigates *⍺*-Syn accumulation and thereby prevents the degeneration of SN DA neurons.

Our previous study demonstrated the upregulation of *Fer2* in the *Drosophila* brain following a brief oxidative insult that triggers the selective loss of DA neurons. A similar pattern was also observed in the *C. elegans* homolog, suggesting that these genes act as stress-response genes that counteract neurotoxins to prevent dopaminergic neurodegeneration (Bou Dib *et al*., 2014). Given the marked functional conservation between invertebrate and vertebrate homologs, we next asked if *Nato3* would be similarly upregulated after neurotoxin exposure. By analyzing *Nato3* mRNA expression in the SN by reverse-transcription quantitative PCR (RT-qPCR), we observed a significant upregulation of *Nato3* immediately following MPTP injection compared to a saline control. *Nato3* levels returned to baseline within one week after five consecutive MPTP injections (Fig. 2j). This rapid reaction is similar to that of the invertebrate homologs (Bou Dib *et al*., 2014), further supporting NATO3’s role as a protective factor against neurotoxic insults.

### Lentiviral-mediated *NATO3* overexpression in patient-derived DA neurons

Human induced pluripotent stem cells (iPSCs) provide access to mDA neurons from PD patients that retain genetic risk variants and offer a tool that circumvents the limited similarity between animal models and humans (Singh Dolt *et al*., 2017). Our findings that *Nato3* overexpression protects DA neurons in mouse PD models prompted us to investigate if overexpression of its highly conserved human homolog, *NATO3* (*FERD3L*), confers neuroprotection in PD patient-derived mDA neurons.

*SNCA* encodes the presynaptic protein α-Syn, the major component of Lewy bodies (Spillantini *et al*., 1997; Baba *et al*., 1998). A53T missense mutation in the *SNCA* gene is associated with autosomal dominant, early-onset PD (Polymeropoulos *et al*., 1997). To investigate the neuroprotective potential of *NATO3*, a human iPSC line derived from a patient carrying the A53T *SNCA* point mutation, and its isogenic mutation-corrected control line were obtained from the National Institute of Neurological Disorders and Stroke (NINDS) Repository. These iPSC lines were differentiated into mDA neurons following the established protocol (Tofoli *et al*., 2019). The differentiation was initiated at day 0 *in vitro* (DIV0) and completed by DIV30 (Fig. 3a). The efficacy of the differentiation into DA neurons was validated through immunostaining for TH. The protocol resulted in the generation of approximately 30% of cells differentiated into TH-positive neurons from both the PD and control iPSCs (Supplementary Figs. 2a and b).

**Figure 3.**
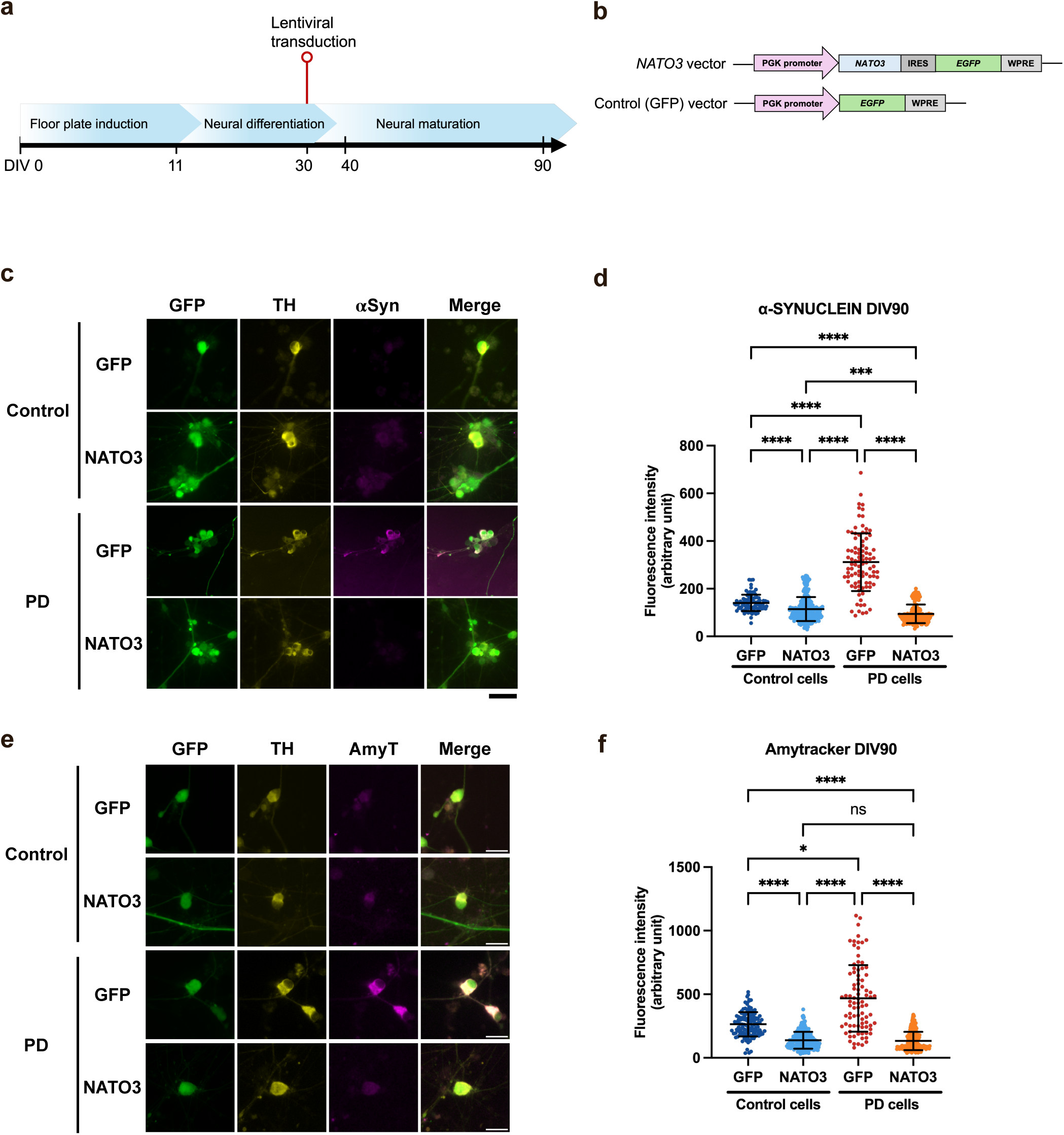
*NATO3* overexpression reduces the abnormal buildup of α-Syn and protein aggregates in DA neurons derived from a PD patient’s iPSCs. **a.** Experimental timeline. **b.** Schematic of the lentiviral vector for bicistronic expression of NATO3 and EGFP from the PGK promoter, and the control vector expressing only EGFP. **c.** Representative images of iPSC-derived DA neurons immunostained for α-Syn, TH, and GFP. PD, cells differentiated from an iPSC line carrying the *SNCA* A53T mutation. Control, mutation-corrected isogenic control cells. NATO3, cells transduced with the NATO3 vector. GFP, cells transduced with the control vector. Scale bar, 50 µm. **d.** Quantification of α-Syn fluorescence intensity within DA neurons at DIV90. **e.** Representative images of iPSC-derived DA neurons stained with Amytracker 630 (AmyT), co-stained for TH and GFP. Scale bar, 20 µm. **f.** Quantification of Amytracker fluorescence intensity within DA neurons at DIV90. In **d** and **f,** scatter plots indicate mean ± SD. Each data point represents the mean fluorescence intensity value from a single TH-positive cell. All data are from N=3 independent differentiation experiments. Statistical comparisons were performed using a Kruskal-Wallis test with a Dunn’s multiple comparisons test. *p<0.05, **p<0.01, ***p<0.001, and ****p<0.0001. ns, not significant.

The A53T mutation increases the stability of α-Syn protein and its propensity to form protofibrils (Conway *et al*., 2000). Evidence suggests that α-Syn misfolding contributes to PD pathogenesis through various cellular dysfunctions, such as mitochondrial and lysosomal impairments and synaptic dysfunctions (Calabresi *et al*., 2023; Park *et al*., 2025). Aligning with this notion, previous studies have shown that α-Syn protein levels are increased in neurons derived from the iPSC lines carrying the *SNCA* A53T mutation compared to the control cells. Additionally, these lines present impaired neurite outgrowth in DA neurons (Kouroupi *et al*., 2017; Zambon *et al*., 2019). To determine if these reported characteristics are present in our culture, we monitored α-Syn levels and neurite length in mDA neurons differentiated from the PD and the isogenic control lines. Immunostaining for α-Syn and TH at DIV40 and DIV90 revealed a significant increase in α-Syn levels and reduced neurite length in the PD DA neurons compared to the control neurons at both time points (Supplementary Fig. 2c and g), consistent with previous reports. These results confirm that our PD line retains key pathological characteristics.

*NATO3* overexpression in the PD and control cell lines was achieved with two distinct lentiviral vectors. The *NATO3-IRES-GFP* vector was designed for the bicistronic expression of both NATO3 and GFP under the control of the murine phosphoglycerate kinase (PGK) promoter, utilizing an Internal Ribosomal Entry Site (IRES). The control vector expressed only GFP under the PGK promoter (Fig. 3b). Since NATO3 is involved in the development of mDA neurons (Ono *et al*., 2010; Nissim-Eliraz *et al*., 2013; Peterson *et al*., 2019), lentiviral transduction was performed on fully differentiated mDA neurons at DIV30 to avoid affecting the iPSC differentiation process. On average, more than 50% of the mDA neurons were transduced, judging from the number of cells co-expressing TH and GFP (Supplementary Fig. 3a). *NATO3* mRNA levels in non-transduced and transduced cells from both cell lines were measured by RT-qPCR. *NATO3* endogenous levels were similar between the control and PD cell lines, with a substantial increase following the NATO3 vector transduction (Supplementary Fig. 3b).

### *NATO3* overexpression prevents aberrant α-Syn overaccumulation and accumulation of misfolded protein in patient-derived DA neurons

Following the verification of our experimental system, we investigated the effect of *NATO3* overexpression in PD neurons and control neurons on various characteristics associated with PD pathology at DIV90, considering that PD pathogenesis is principally age-dependent. To account for any nonspecific effects of lentiviral transduction, we compared the phenotypes between the cells transduced with the NATO3 vector and those with the GFP vector. Using GFP marker and anti-TH staining, the immunostaining analysis focused specifically on transduced DA neurons. We first examined α-Syn levels by immunohistochemistry. Elevated α-Syn levels in PD cells compared to control cells remained unaltered by the transduction of the control vector. Whereas the NATO3 vector transduction substantially reduced α-Syn levels in PD cells to the levels of the control cells (Figs. 3c and d).

The presence of protein aggregates, notably Lewy bodies and the aggregates of the microtubule-associated protein Tau, is a pathological hallmark of PD (Goedert *et al*., 2013). We next monitored protein aggregates in iPSC-derived mDA neurons using the Amytracker 630 marker, which stains repetitive arrangements of β-sheet structures typically found in amyloids (Morten *et al*., 2022). Amytracker staining revealed a significant accumulation of aggregates in PD neurons compared to control neurons. *NATO3* overexpression reversed this increase, as well as reduced the Amytracker levels in the control cells (Figs. 3e and f). Taken together, these results indicate that *NATO3* overexpression reduces misfolded protein accumulation and counteracts α-Syn overaccumulation, both linked to the *SNCA* A53T mutation.

### *NATO3* overexpression restores autophagic function and preserves mitochondria in patient-derived DA neurons

The A53T point mutation and resulting intracellular accumulation of α-Syn have been shown to disrupt the autophagy-lysosomal pathway (Abeliovich & Gitler, 2016). We therefore investigated the status of autophagy in iPSC-derived DA neurons by staining for the selective autophagy receptor P62. We observed a significant increase in P62 levels in PD neurons compared to the control neurons, indicating the defective autophagic flux in PD DA neurons. *NATO3* overexpression in control neurons increased P62 levels compared to the expression of the GFP vector, suggesting that NATO3 accumulation affects basal autophagy levels. However, in PD neurons, *NATO3* overexpression restored P62 levels to control levels (Figs. 4a and b). These results suggest that *NATO3* overexpression can stimulate autophagic flux in pathological conditions, thereby restoring the defective autophagy. This finding is strikingly congruent with our *in vivo* data, which show that loss of *Nato3* leads to autophagic blockage, while its overexpression stimulates it in mice (Figs. 2b-g).

**Figure 4.**
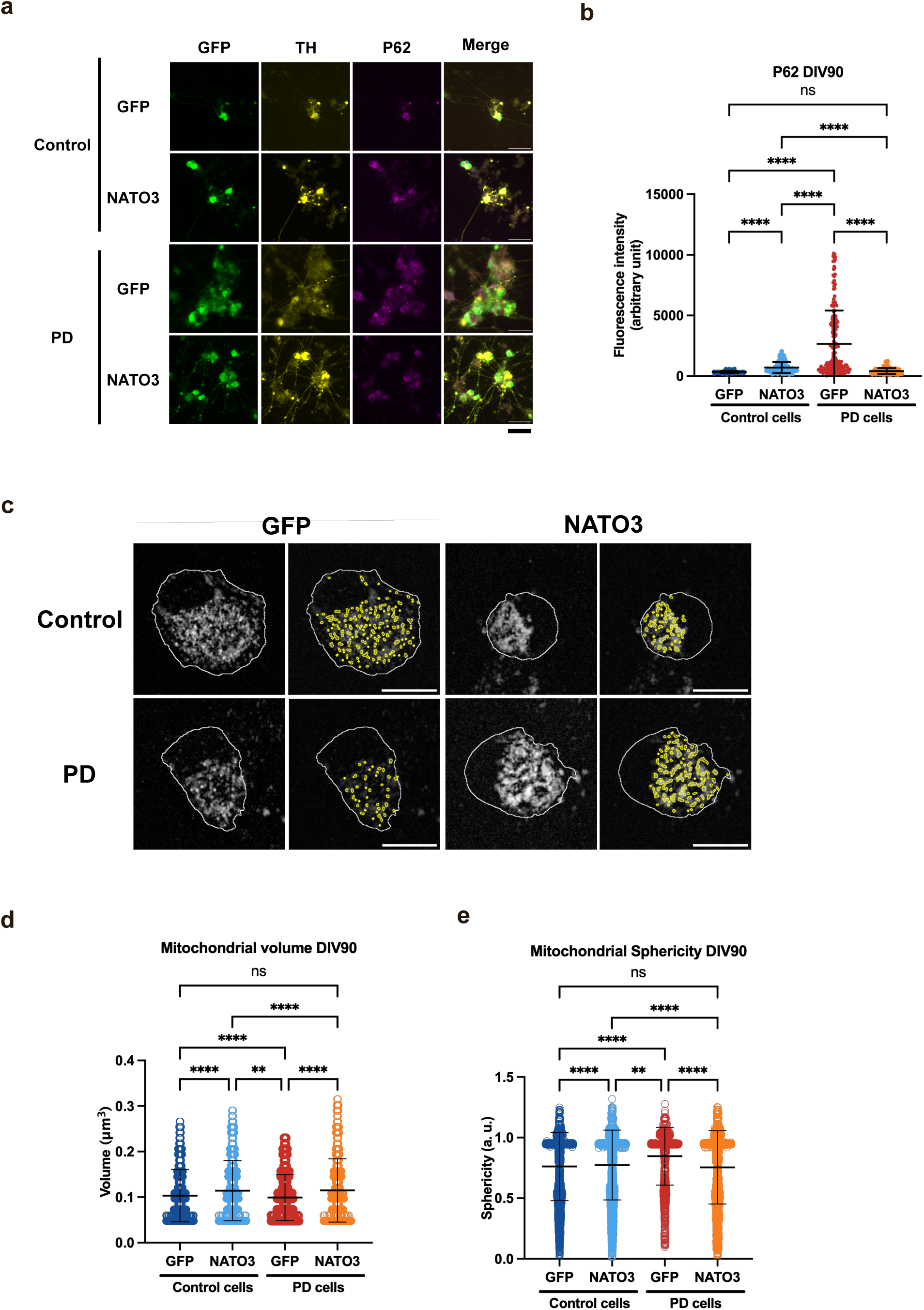
*NATO3* overexpression improves autophagic flux and restores aberrant mitochondrial morphology in DA neurons derived from a PD patient. **a.** Representative images of iPSC-derived DA neurons immunostained for P62, TH, and GFP. Control and PD cell lines, transduced with the NATO3 vector (NATO3) or the control vector (GFP). Scale bar, 50 µm. **b.** Quantification of P62 fluorescence intensity within DA neurons at DIV90. **c.** Representative high-magnification images of iPSC-derived DA neurons immunostained for TOM20, an outer mitochondrial membrane protein (gray). The cell body of a TH-positive cell is outlined in white. Mitochondria are outlined in yellow, a feature generated by the Mitochondria Analyzer plugin in Fiji. Scale bar, 10 µm. **d and e.** Quantification of mitochondrial volume (**d**) and sphericity (**e**) marked by TOM20 in DA neurons at DIV90. In **b, d,** and **e,** scatter plots indicate mean ± SD. Each data point represents the value from a single TH-positive cell. All data are from N=3 independent differentiation experiments. Statistical comparisons were performed using a Kruskal-Wallis test with a Dunn’s multiple comparisons test. **p<0.01 and ****p<0.0001. ns, not significant.

Defective autophagy-lysosomal pathway is tightly linked to mitochondrial impairments, characterized by disruptions in morphology, dynamics, and functions, which are pathological hallmarks of PD (Henrich *et al*., 2023). Abnormal changes in mitochondrial shape have been specifically associated with α-Syn oligomers and aggregates (Xie & Chung, 2012a; Plotegher *et al*., 2014), indicating a mechanistic link between α-Syn and mitochondrial morphology in PD. We therefore assessed mitochondrial morphology in iPSC-derived DA neurons by immunostaining for TOM20, a mitochondrial outer membrane protein. Compared to control cells, PD neurons exhibited mitochondrial fragmentation, characterized by decreased mitochondrial volume and increased sphericity (Figs. 4c-e). This suggests a disruption in the fission-fusion dynamics, which are essential for maintaining a healthy mitochondrial network. While fission is required for the removal of damaged mitochondria by autophagy (Twig & Shirihai, 2011; Pickrell & Youle, 2015), enhanced fission and decreased fusion can lead to a failure in this process. We found that overexpression of *NATO3* restored both parameters of mitochondrial morphology (Figs. 4c-e). These results indicate a role for *NATO3* in maintaining mitochondrial health, as previously shown in mice (Miozzo *et al*., 2022). This effect is, at least in part, likely mediated by the role of *NATO3* in stimulating the autophagy-lysosomal pathway.

### *NATO3* overexpression reverses aberrant neurite morphology in patient-derived DA neurons

Axonal degeneration has a central role in PD pathogenesis, highlighted by the decreased number of DA neuron projections in the striatum, which leads to its motor symptoms (Hornykiewicz, 1998; Cheng *et al*., 2010). Dystrophic neurites have been identified in the brains of α-Syn A53T patients (Duda *et al*., 2002). This trait has also been observed in DA neurons derived from the iPSCs carrying the same mutation (Kouroupi *et al*., 2017; Czaniecki *et al*., 2019).

We analyzed the length and complexity of neurites in iPSC-derived DA neurons at DIV40 and 90 to assess the effect of the mutation and *NATO3* overexpression in the development and maintenance of neurites. We observed the pathological changes in neurites in PD cells, congruent with the previous reports. At both time points, the length of the neurites was reduced in PD neurons compared to the control neurons. *NATO3* overexpression extended the neurites of PD neurons (Figs. 5a-c). Sholl analysis, the commonly used method to quantify the complexity of neuronal projections (Binley *et al*., 2014), revealed reduced complexity in PD DA neuron projections at both DIV40 and 90. *NATO3* overexpression reversed this trend, increasing the neurite complexity of the PD cells to the levels of the control cells at both time points (Figs. 5a, d-f, and Supplementary Table 1). These results reveal that *NATO3* overexpression counteracts the detrimental effect of the α-Syn A53T mutation on neurite development and maintenance. Given the critical role of neurite length in synaptic connectivity and signal transmission, it is likely that *NATO3* overexpression restores neuronal functions impaired by the α-Syn A53T mutation.

**Figure 5.**
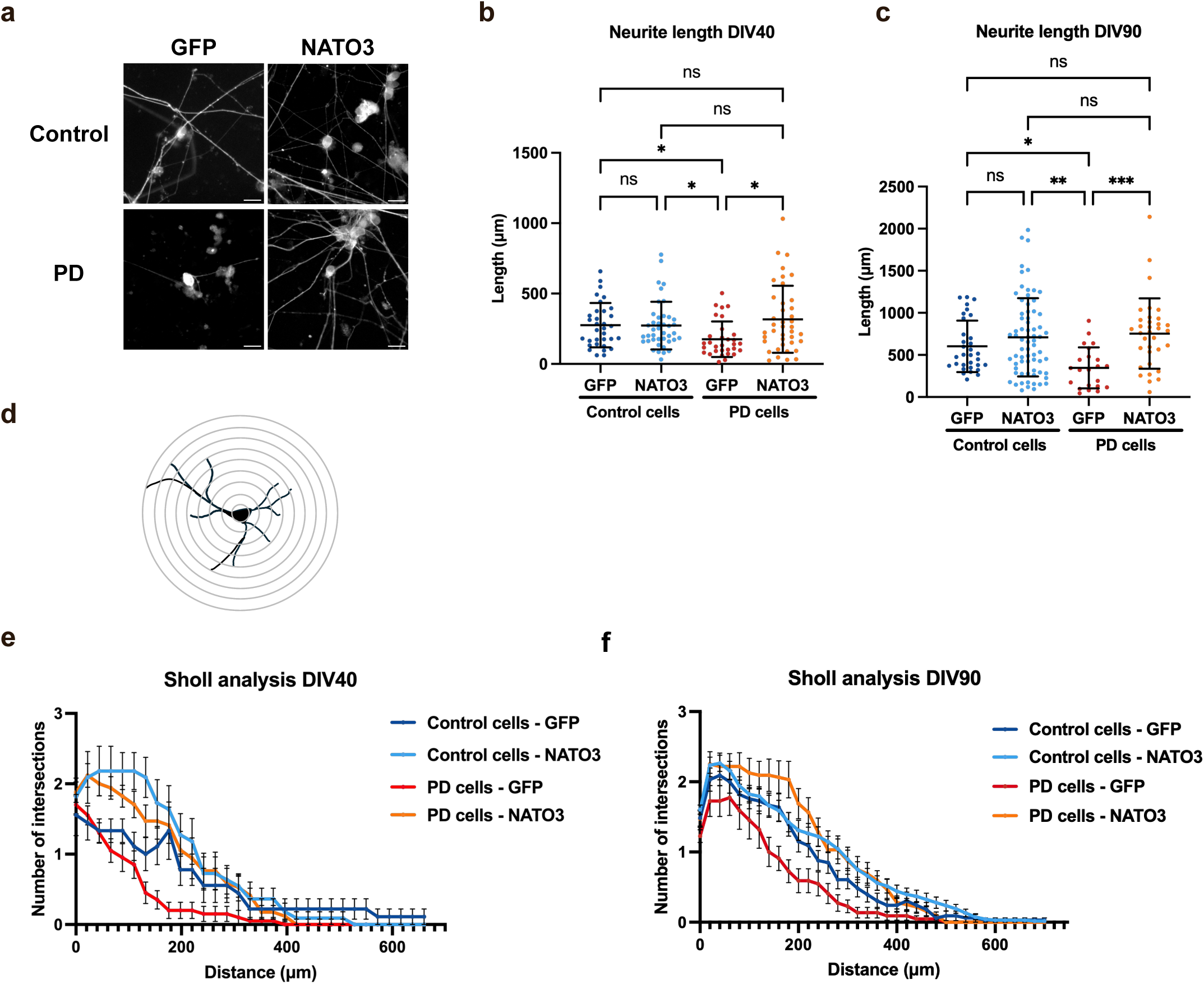
*NATO3* overexpression restores aberrant neurite length and complexity in DA neurons derived from a PD patient. **a.** Representative images of iPSC-derived DA neurons immunostained for TH at DIV90. Control and PD cell lines, transduced with either the NATO3 vector (NATO3) or the control vector (GFP). Scale bar, 50 µm. **b and c**. Neurite length of TH-positive cells analyzed at DIV40 (**b**) and DIV90 (**c**). Mean ± SD. Each data point represents the value from a single TH-positive cell. N=3 independent differentiation experiments. Statistical comparisons were performed using a Kruskal-Wallis test with a Dunn’s multiple comparisons test. *p<0.05, **p<0.01, and ***p<0.001. ns, not significant. **d.** Diagram illustrating Sholl analysis, a quantitative method for assessing the complexity and branching patterns of neuronal processes by measuring the number of intersections with concentric circles. **e and f.** Results of Sholl analysis of DA neurons at DIV40 (**e**) and DIV90 (**f**). The x-axis indicates the distance from the center of the cell body, and the y-axis indicates the number of neurites crossing the given concentric circle. Mean ± SD. N=3 independent differentiation experiments. Statistical comparisons were performed using a two-way ANOVA followed by Tukey’s multiple comparisons test, and the results are displayed in Supplementary Table 1.

## Discussion

Neurodegeneration presents a persistent challenge in finding an effective therapeutic solution due to its complex and multi-factorial root causes. Our findings demonstrate that an overexpression of a single transcription factor can mitigate pathological characteristics that lead to degeneration of mDA neurons in both *in vivo* and human *in vitro* models.

To examine the neuroprotective potential of NATO3 in an *in vitro* model of PD, we opted for an iPSC line derived from a PD patient harboring the α-Syn A53T mutation associated with autosomal dominant PD. The A53T mutation in α-Syn protein promotes β-sheets formation, leading to a faster rate of fibril formation as a toxic gain of function, which contributes to the early onset of familial PD (Conway *et al*., 1998; Bertoncini *et al*., 2005; Lashuel *et al*., 2013). The α-Syn A53T cell lines used in prior studies consistently displayed PD-linked phenotypes across different cell lines and laboratories, despite batch effects typically associated with iPSC experiments. These include elevated levels of α-Syn, early protein aggregates, mitochondrial dysfunction, and autophagy defects (Dettmer *et al*., 2015; Mazzulli *et al*., 2016; Kouroupi *et al*., 2017; Prots *et al*., 2018; Stykel *et al*., 2018; Czaniecki *et al*., 2019; Zambon *et al*., 2019; Snowden *et al*., 2020; Diao *et al*., 2021). These phenotypes were also observed in our differentiated mDA neurons derived from the α-Syn A53T PD line when compared to the mutation-corrected isogenic control line.

Overexpression of *Nato3* (mouse) or *NATO3* (human) reduces the aberrant accumulation of α-Syn protein in mice and in patient-derived mDA neurons, respectively. This effect is likely attributed to the role of *Nato3* homologs in stimulating autophagic flux. While the precise molecular mechanism linking how *Nato3* regulates the autophagy-lysosomal pathway remains to be elucidated, its involvement in autophagy regulation has been indirectly supported by a genome-wide CRISPR screen (Takahashi *et al*., 2019). *Nato3* regulates a genetic feed-forward loop involving key DA neuron developmental transcription factors, including LMX1B, FOXA2, and SHH (Nissim-Eliraz *et al*., 2013; Peterson *et al*., 2019; Miozzo *et al*., 2022). Because it has been shown that Lmx1b regulates autophagy-lysosomal function in mDA neurons (Laguna *et al*., 2015; Jiménez-Moreno *et al*., 2023), the ability of Nato3 to stimulate autophagic flux may be mediated, at least in part, through this transcriptional network.

Amytracker signal levels are increased in the patient-derived mDA neurons, indicating an aberrant accumulation of amyloid-like β-sheet structures (Mahul-Mellier *et al*., 2020). The increase in Amytracker fluorescence likely corresponds to the accumulation of A53T mutant α-Syn undergoing oligomerization and forming β-sheets. However, alternative misfolded proteins undergoing similar processes might also be labelled by Amytacker. While α-Syn is the primary component found in Lewy bodies within the SN of PD patients, other proteins contribute to their composition (Goldman *et al*., 1983; Kawamoto *et al*., 2002; Wakabayashi *et al*., 2007). *NATO3* overexpression mitigates overaccumulation of both α-Syn and Amytracker signals, consistent with the finding that NATO3 enhances autophagic activity that facilitates clearance of misfolded proteins.

Mitochondria are dynamic organelles that continuously undergo fusion and fission to respond to cellular demands. Fusion generates an interconnected mitochondrial network, facilitating inter-organelle communication, matrix content diffusion, mitochondrial DNA repair, and metabolite distribution among mitochondria. Fission creates smaller mitochondria that tend to generate more ROS, aids equal mitochondrial segregation during cell division, and enhances the distribution of mitochondria along cytoskeletal tracks. Fission also facilitates the isolation of damaged mitochondrial segments to promote their autophagy. Therefore, mitochondrial remodeling is directly linked to the maintenance of mitochondrial functions, and its dysregulation has been reported in neurodegenerative diseases (Youle & van der Bliek, 2012; Henrich *et al*., 2023; Brown *et al*., 2025). Our PD cell line presented smaller and more spherical mitochondria, suggesting a fragmentation in the mitochondrial network. This observation aligns with prior studies on human iPSC and mouse models with the A53T mutation, which reported an increase in mitochondrial fission (Xie & Chung, 2012b; Zambon *et al*., 2019). Our data are also consistent with the reports that α-Syn directly associates with mitochondrial membranes, leading to mitochondrial damage, including mitochondrial fragmentation and complex I inhibition (Nakamura *et al*., 2011; Serdiuk *et al*., 2025).

The mitochondrial dysmorphia and fragmentation observed in patient-derived DA neurons are restored by *NATO3* gene delivery. This is congruent with the role of NATO3 in stimulating autophagic flux. The selective dopaminergic neurotoxin MPTP primarily targets mitochondrial electron transport chain (ETC) complex I. This results in ATP deficits and an increase in ROS production, leading to the degeneration of DA neurons (AlShimemeri *et al*., 2021). Our finding that Nato3 overexpression in mice counteracts MPTP-induced DA neurons loss suggests that stimulation of autophagy-lysosomal function by Nato3 can ameliorate mitochondrial impairments. However, this does not exclude the possibility that Nato3 may also more directly regulate mitochondrial dynamics, quality, or quality control.

*NATO3* overexpression counteracts the detrimental effect of the α-Syn A53T mutation in the development and maintenance of neurites in DA neurons. This finding suggests that NATO3’s role is not limited to stimulating the autophagy-lysosomal pathway. Interestingly, prior research has shown that overexpression of *NATO3* alone can direct human embryonic stem cells (hESCs) to differentiate into neuron-like cells possessing neurites (Joung *et al*., 2023). This may reflect the developmental role of NATO3, since it is downstream of FOXA2 and suppresses HES-1. Inactivation of HES-1 induces neurite outgrowth mediated by NGF signaling (Ström *et al*., 1997). Additionally, a systems-level transcriptome analysis identified NATO3 as a highly ranked gene for its role in the dynamic regulation of neural differentiation of both mouse ESCs and human iPSCs (Ando *et al*., 2015). These prior findings and our results taken together highlight the relevance of NATO3 for stem cell therapy.

*NATO3* is a potential target of the sterol regulatory element binding protein 1 (SREBP1), encoded by the *SREBF1* gene (Toledo *et al*., 2020). Notably, a polymorphism in the *SREBF1* gene is linked to an increased risk of PD (Do *et al*., 2011; Shulman *et al*., 2014). Additionally, *SREBF1* has been shown to regulate mitophagy (Ivatt *et al*., 2014). These prior findings suggest that dysregulation of *NATO3* expression could contribute to PD pathogenesis. Furthermore, genome-wide DNA methylation analysis has found that CpG sites in the *NATO3* gene are significantly hypo-methylated in the SN and the dorsal motor nucleus of the vagus (DMV) in PD patients compared to controls (Young *et al*., 2019). This epigenetic change may either suggest a role in pathogenesis or indicate that *NATO3* expression is induced to cope with pathogenic processes. The latter possibility is supported by our observation that MPTP treatment upregulates *Nato3* expression.

Matched by the crucial role of mDA neuron developmental genes for the survival of these neurons during aging, gain-of-function studies—such as the overexpression of *Engrailed* (Alvarez-Fischer *et al*., 2011), the combined overexpression of *Nurr1* and *FoxA2* (Oh *et al*., 2015), and the use of NURR1 agonist treatments (Kim *et al*., 2023)— have demonstrated neuroprotective effects in mouse PD models. Adding *Nato3* to this list of potential therapeutic targets enriches the options for gene and stem cell therapy approaches aimed at disease modification.

A limitation of our mouse experiments is that *Nato3* overexpression was conducted before the induction of neuronal loss with MPTP or together with the overexpression of α-Syn. While this approach was chosen due to the rapid action of MPTP and to avoid two consecutive stereotactic injections, it means that our findings represent a preventive effect rather than an intervention during disease progression. Given the current lack of robust early disease biomarkers for PD, future work should focus on determining the specific stage at which NATO3 expression can effectively intervene in disease progression.

## Materials and Methods

### Mouse strains

*Nato3* cKO mouse strain, which is homozygous for the *Nato3-LoxP* allele and harbors one copy of the *DAT-Cre* transgene (*Nato3^DAT-Cre^*), is described in (Miozzo *et al*., 2022). C57BL/6 was used as a wild-type strain. Both male and female mice were used in all the experiments. Mice were maintained in rooms with controlled 12h light/dark cycles, temperature between 23 and 24 °C, and humidity of 47–61%, with food and water provided *ad libitum*. Mice were housed at a maximum of five animals per cage in individually ventilated cages.

### Study approval

All animal experiments were conducted in accordance with the Institutional Animal Care and Use Committee of the University of Geneva and with permission of the cantonal authorities (License No. GE29B 33210 and GE/266C 34952).

### Behavioral tests

All behavioral tests were performed during the light cycle. Mice were allowed to habituate to the behavioral room for at least 45 min before each test. Behavioral equipment was cleaned with 70% ethanol after each test session to avoid olfactory cues. The pole test was performed as previously described (Matsuura *et al*., 1997) with minor modifications. Mice were first trained 4–6 times by placing the animal head-down on top of a vertical pole (diameter: 1 cm, height: 55 cm) and letting them descend. Then, animals were trained 3–4 times in the regular turning and descending procedure, by placing the animal head-up on top of the pole. For the actual test, mice performed nine trials (three trials per day for three consecutive days). Each trial was recorded with a video camera, with an interval of at least 5 min between trials. The time to orient downward (T-turn) and the total time to turn and descend the pole (T-total) were measured, with a maximum duration of 30 seconds. If the mouse was not able to turn downward and instead dropped from the pole in a lateral body position, a value of 30 seconds was assigned. The average of the nine trials was used as the final score.

### Viral vectors

The lentiviral vector for NATO3 overexpression was based on an expression cassette driven by the mouse PGK1 promoter and the woodchuck posttranscriptional regulatory element (WPRE). The human NATO3 (FERD3L) (NM_152898.2) coding sequence tagged with an N-terminal HA epitope tag was subcloned in front of an internal ribosome entry site (IRES) sequence followed by the EGFP coding sequence, and the woodchuck posttranscriptional regulatory element (WPRE). This synthetic construct was custom-made by Eurofins and cloned into a lentiviral transfer plasmid backbone. The control vector, which also used the same backbone, contained the EGFP coding sequence under the PGK promoter and the WPRE. Both vectors were packaged into self-inactivating particles pseudotyped with the vesicular stomatitis virus G protein (VSV-G) envelope. Lentiviral titer was estimated by P24 protein concentration measured using a p24 ELISA (ZeptoMetrix, NY, USA), with the conversion 1pg P24 = 50 TUs.

The pAAV construct for human *⍺*-Syn expression (AAV7-syn1-*⍺*Syn-WPRE) was based on the neuron-specific *synapsin 1* (syn1) promoter and the WPRE sequence. The vector was designed to encode the human wild-type *⍺*-Syn coding sequence (NM_000345), as described in (Oliveras-Salvá *et al*., 2013). The corresponding control AAV vector (AAV7-syn1-empty-WPRE) lacked the *⍺*-Syn insert. Both constructs were packaged into serotype 7 AAV particles using suspension-adapted HEKExpress™ cells and purified using an AAVX column for affinity chromatography, similarly to the method described in (Gaudry *et al*., 2024). The construct for expressing *Nato3* (AAV6-pgk-*mNato3*-WPRE) included the mouse PGK1 promoter, the beta globin intron, the mouse *Nato3* coding sequence (NM_033522.2), and the WPRE enhancer, and was similarly packaged into serotype 6 AAV particles. The genomic titer (VG) of the AAV vector suspensions was determined by digital PCR using a 2-plex QIAcuity apparatus (QIAGEN). All vector productions were performed at the EPFL Bertarelli Foundation Gene Therapy Platform.

### Stereotaxic injection

Mice were anesthetized with isoflurane before receiving a unilateral injection in the substantia nigra (right hemisphere). A total of 2 μL of vector suspension diluted in PBS was injected per mouse. The titer of the AAV7-syn1-Syn-WPRE vector suspension expressing the wild-type human α-synuclein protein was 3.4 × 10^13^ VG/mL. The titer of the AAV7-syn1-empty-WPRE vector suspension was 2.6 × 10^13^ VG/mL. The injected vector dose was set at 1.5 × 10^9^ VG for the AAV-αSyn and the control AAV vectors. For *Nato3* overexpression, the titer of the AAV6-pgk-Nato3-WPRE vector suspension was 5.4 × 10^11^ VG/mL. The injected vector dose was set at 1.08 × 10^9^ VG. In the control group, animals were injected with the same volume of PBS. The injections were performed using a standard stereotaxic procedure (Landeck *et al*., 2021) with the following coordinates relative to Bregma: −3.0 mm (anteroposterior), −1.3 mm (mediolateral), and −4.7 mm (dorsoventral).

### MPTP administration

Mice received intraperitoneal (IP) injections of MPTP hydrochloride (Sigma, Ref#5063820001) at a dose of 20mg/kg body weight. MPTP was dissolved in saline (0.9% NaCl) and administered for 5 consecutive days. Control mice received saline injections only.

### Brain collection and sectioning

At selected time points, mice were anesthetized with pentobarbital before intracardiac perfusion. Mice were first perfused with 0.2 % heparin (Sigma-Aldrich #H3149-25KU) in PBS in the left ventricle of the heart to prevent blood coagulation, followed by ice-cold 10% formalin solution (Sigma HT501128-4L). Subsequently, brains were collected and post-fixed in 4% paraformaldehyde (PFA, Thermo Fisher Scientific #28906) for three nights at 4°C. The fixed brains were then stored in PBS at 4°C. For sectioning, brains were embedded in 6% low-melting agarose (Thermo Fisher Scientific #R0801). Using a vibratome, 40 μm-thick coronal sections were cut through the striatum and the midbrain. The sections were collected in PBS and stored at 4°C in 24-well plates for subsequent analysis.

### iPSC culture and differentiation into midbrain DA neurons

Human iPSC lines were obtained from the NINDS Repository at the Coriell Institute for Medical Research: a PD patient line harboring the *SNCA* A53T point mutation (cat. no. ND50086) and the mutation corrected isogenic control line (cat. no. ND50085). The iPSCs were maintained on Geltrex (A1413302, Thermo Fisher Scientific) in StemFlex basal medium supplemented with 10% StemFlex supplement (10x stock) (A3349401, Thermo Fisher Scientific) and passaged using Versene solution (BE17-711E, Lonza). All media contained 1% penicillin-streptomycin (100x) (15140-122, Thermo Fisher Scientific). Cells were cultured at 37°C in a humidified atmosphere containing 5% CO_2_.

Differentiation of iPSCs into midbrain DA neurons was performed as previously described (Tofoli *et al*., 2019). Briefly, iPSCs were seeded onto Geltrex-coated six-well plates with daily medium changes. When the cultures reached >80% confluency (designated as day 0 *in vitro*; DIV), StemFlex media was replaced with KSR medium, composed of KnockOut DMEM (10829-018, Thermo Fisher Scientific), 1% MEM NEAA (100x) (11140035, Thermo Fisher Scientific), 1% Glutamax (100x) (35050038, Thermo Fisher Scientific), 0.1% β-Mercaptoethanol (1000x) (21985-023, Thermo Fisher Scientific) and 15% KnockOut Serum Replacement (10828028, Thermo Fisher Scientific), supplemented with 100 nM LDN193189 (130-103-925, Stem Macs) and 10 µM SB431542 (1614/10, Tocris Bioscience).

From DIV1 to DIV4, the medium was changed daily On DIV1 and 2, cultures were additionally supplemented with 100 ng/ml Sonic Hedgehog (SHH) C25II (464-SH-200, R&D Systems), 2 µM Purmorphamine (540220-5MG, Sigma-Aldrich), and 100 ng/ml Fibroblast growth factor 8 (FGF-8b; 423-F8-025, R&D systems). On DIV3 and DIV4, the medium was further supplemented with 3 µM CHIR99021 (361571, Milipore).

Starting at DIV5, SB431542 supplementation was discontinued, and KSR medium was gradually replaced with N2 medium, composed of DMEM F12 (11320033, Thermo Fisher Scientific), Glutamax (35050038, Thermo Fisher Scientific), and N2 supplement (17502048, Thermo Fisher Scientific). The proportion of N2 medium was increased every two days: 25% (DIV5), 50% (DIV7), and 75% (DIV9). From DIV7 onward, the supplementation of LDN193189 (100 nM) and CHIR99021 (3 µM) was maintained, while Purmorphamine and FGF8 were discontinued.

On DIV11, medium was changed to neuronal maturation medium, consisting of Neurobasal Medium (21103049, Thermo Fisher Scientific), 2% B27 supplement (50x) (17504044, Thermo Fisher Scientific) and 1% Glutamax (100x) (35050038, Thermo Fisher Scientific), supplemented with 20 ng/ml Brain Derived Neurotrophic Factor (BDNF; 248-BDB-010, R&D Systems), 200 µM ascorbic acid (A4544-100G, Sigma-Aldrich), 20 ng/ml Glial Derived Neurotrophic Factor (GDNF; 212-GD-050, R&D Systems), 1 ng/ml transforming growth factor beta 3 (TGFb3; 243-b3-002, R&D Systems), 250 µM dibutyryl cAMP (db cAMP, D0260-100MG, Sigma-Aldrich) and 10 µM DAPT (2634/10, Tocris Bioscience). Additionally, 3 µM CHIR99021 was supplemented on DIV11 and DIV12.

On DIV20, cells were dissociated using Accutase (561527, BD Biosciences) and reseeded at a density of 2 × 10^5^ cells/cm^2^ onto 24-well plates coated with Geltrex. 10 µM Y27632 (ROCK inhibitor; SCM075, Sigma-Aldrich) was added to the medium after passages. From DIV21 onward, medium was changed every two days until mDA neurons reached the desired maturation stage.

### Lentiviral transduction

iPSC-derived DA neurons were passaged on DIV20 and plated at a density of 2 × 10^5^ cells/cm^2^ onto sterilized, Geltrex-coated coverslips placed in 24-well plates. Lentiviral transduction was performed on DIV30 using the same amount of P24 corresponding to the Multiplicity of Infection (MOI) of 5. The following day, the medium was replaced, and neurons were maintained in neuronal maturation medium until the designated experimental time points.

### RT-qPCR

RT-qPCR on mouse midbrain tissues was performed as described previously (Miozzo *et al*., 2022), with minor modifications. Total RNA was isolated from the dissected SN using TRIzol Reagent (Life Technologies) according to the manufacturer’s instructions. Reverse transcription was performed from 80 ng of RNA. First, RNA samples were incubated with double-strand-specific DNase at 37°C for 2 min to remove genomic DNA. Then, RNA samples were reverse transcribed with Maxima cDNA H Minus Synthesis Master Mix (M1682, Thermo Fisher Scientific) containing oligo(dT)_18_ and random hexamer primers for 10 min at 25°C, followed by 15 min at 50°C and 5 min at 85°C. Quantitative PCR was carried out with PowerTrack™ SYBR™ Green Master Mix (C14512, Applied Biosystems) on a QuantStudio 5 PCR instrument (Thermo Fisher). *Nato3* primer efficiency and housekeeping gene *Gapdh* were validated prior to the experiment and negative controls were included. *Nato3* expression levels were normalized to *Gapdh* levels. The normalized *Nato3* mRNA levels in the MPTP-treated groups were then compared to the mean expression levels in the NaCl-treated animals (D14). For RT-qPCR on iPSC-derived DA neurons, total RNA was extracted with TRIzol Reagent and treated with Turbo DNase-free kit. DNA removal and reverse transcription was performed using the Maxima H Minus cDNA Synthesis Master Mix with dsDNase (M1682, Thermo Fisher Scientific) and oligo-dT primers, following the manufacturer’s protocol. Relative expression of human *NATO3* was normalized to the expression of housekeeping genes *GADPH*, *ACTB*, and *TBP*, using a multiple reference genes model (Hellemans *et al*., 2007). Quantitative PCR was performed on a QuantStudio 5 PCR system using the PowerTrack SYBR Green Master Mix (C14512, Applied Biosystems). All experiments were carried out in triplicate. Primers were designed with the Primer-BLAST online tool, and the primer sequence information is presented in the table below.

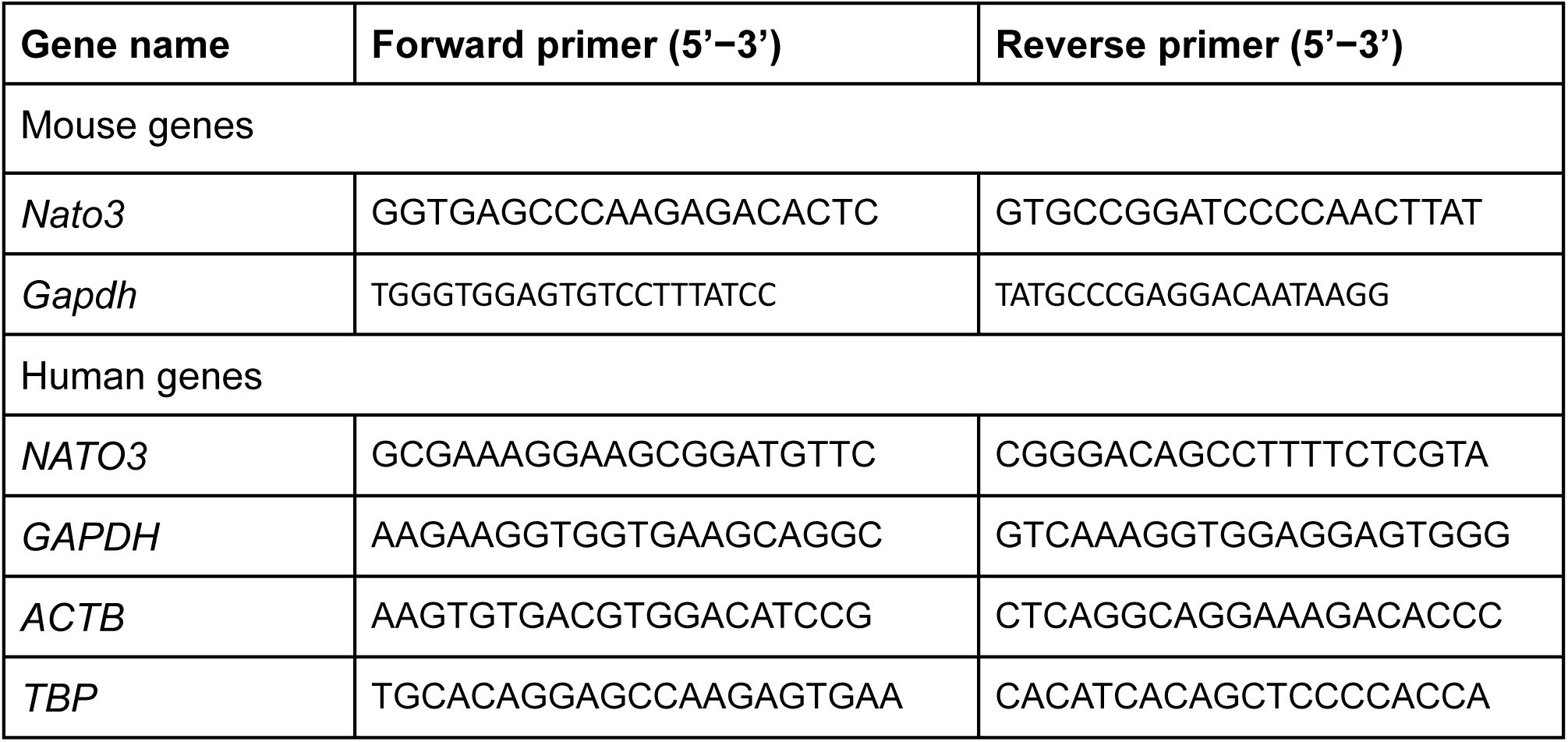

### Immunohistochemistry

Immunostaining of mouse brain tissues was performed on free-floating sections. Sections were first incubated in blocking solution (4% bovine serum albumin (BSA), 4% normal goat serum (NGS), 0.1% Triton X-100 in PBS) for 30 min at RT, followed by incubation with the primary antibody (diluted in 0.1% Tween 20, 4% NGS in PBS) for 48 h at 4°C. Thereafter, samples were washed twice for 30 min each in washing buffer (0.1% Tween 20 in PBS) and incubated with the corresponding secondary antibodies (diluted in 0.1% Tween 20, 4% NGS in PBS) overnight at 4°C in the dark. After two washes, nuclei were counterstained with DAPI (5 µM, Abcam #ab228549) for 30 min at RT. After a 30min final wash, sections were mounted on microscope slides with Vectashield mounting medium (Vectashield PLUS (Vectorlabs, Cat. # H-2000)).

Cells cultured on Geltrex-coated coverslips were fixed with 4% PFA (Thermo Fisher Scientific #28906) for 15 min at RT and washed three times for 10 min with PBS. Samples were then incubated in a permeabilization-blocking buffer (2% NGS, 0.1% Triton X100 in PBS) for 30 min at RT and incubated with primary antibodies (diluted in 2% NGS, 0.1% Tween 20 in PBS) overnight at 4°C. Following the primary antibody incubation, samples were washed with 0.05% Tween-PBS three times for 30 min each, then incubated with the corresponding secondary antibodies (diluted in 2% NGS, 0.1 % Tween 20 in PBS) for 2 h at RT. After three washes of 30 min each in 0.05% Tween-PBS and a final wash of 30 min with PBS, samples were mounted using Vectashield Plus with DAPI (Vectorlabs, Cat. # H-2000). Amytracker staining was performed by incubating the cells for 30 min with Amytracker 630 (1:1000) (Ebba Biotech), followed by three 30-min washes with PBS before mounting. Antibodies and dilutions used in the study are listed in the table below.

| Primary antibodies |  |  |  |  |
| --- | --- | --- | --- | --- |
| Antigen/Epitope name | Host species | Catalog number | Provider | Dilution |
| TH | Rabbit | AB152 | Merck /Sigma-Aldrich | 1:1000 |
| TH | Mouse | 22941 | Immunostar | 1:1000 |
| P62 (SQSTM1) | Mouse | H00008878-M01 | Abnova | 1:500/ 1:1000 |
| TUJ1 | Chicken | TUJ7947980 | AVES | 1:500 |
| α-SYNUCLEIN | Mouse | 610787 | BD Biosciences | 1:500/ 1:1000 |
| TOM20 | Mouse | sc-17764 | Santa Cruz Biotech | 1:500 |

| Secondary antibodies |  |  |  |  |
| --- | --- | --- | --- | --- |
| Fluorophore | Host/<br>Target<br>species | Catalog number | Provider | Dilution |
| Alexa Fluor 633 | Goat anti-Rabbit | A-21070 | Thermo Fisher | 1:500 |
| Alexa Fluor 568 | Goat anti-Rabbit | A-11036 | Thermo Fisher | 1:500 |
| Alexa Fluor 488 | Goat anti-Rabbit | A-11008 | Thermo Fisher | 1:500/ 1:1000 |
| Alexa Fluor 633 | Goat anti-Mouse | A-21050 | Thermo Fisher | 1:500 |
| Alexa Fluor 568 | Goat anti-Mouse | A-11004 | Thermo Fisher | 1:500 |
| Alexa Fluor 488 | Goat anti-Mouse | A-11001 | Thermo Fisher | 1:500 |
| Alexa Fluor 568 | Goat anti-Chicken | A-11041 | Thermo Fisher | 1:500 |

### Microscopy and image analysis

Immunofluorescence images of mouse brain sections were acquired using a Nikon AX confocal microscope, except for the images shown in Fig. 2b, which were obtained using a Nikon Ti / CSU-W1 Spinning Disc Confocal microscope with a 63x air objective. Fluorescence intensity analyses were conducted using Fiji/ ImageJ software (v. 1.54f and 1.54p). Quantification of TH-positive DA neurons in the midbrain was conducted using QuPath software (Bankhead *et al*., 2017) (https://qupath.github.io/) following the method described previously (Berman *et al*., 2011) with minor modifications. Serial 40-µm coronal sections spanning the entire SN were collected using a vibratome. Sections were analyzed at every fourth interval (approximately 12 to 15 sections per animal) with TH immunostaining. Using the polygonal tool of QuPath, SN were delimited and TH-positive DNs were selected based on shape and size parameters. All TH-positive neurons within each physical section, encompassing the full dorsal–ventral and lateral extent of the SN, were counted separately for each hemisphere. Counts from all analyzed sections were summed, and the average number of TH-positive neurons per section was calculated. This value was multiplied by four to estimate the total number of TH-positive neurons per hemisphere for each animal. Fluorescence intensity analyses were conducted using Fiji/ ImageJ software (v. 1.54f and 1.54p). P62 and α-synuclein fluorescence were evaluated within TH-positive neurons that were delimited using the freehand selections tool of Fiji. Then, each TH area was added to the ROI manager. Following an 8-bit conversion of images, P62 and α-synuclein levels were measured as mean grey values within dopaminergic cells across 10 different pictures throughout the SN per hemisphere.

iPSC-derived neurons were imaged using a Leica DM5000 fluorescence microscope at 20x magnification, except for mitochondrial analysis. Image analyses were conducted using Fiji/ ImageJ software (v. 1.54f and 1.54p). Fluorescence intensity was measured within regions of interest (ROI) defined around the neuronal soma of DA neurons identified by TH immunoreactivity. In the case of lentiviral vector-transduced neurons, TH immunoreactivity and co-localizing GFP signal were used to define ROI. Mean fluorescence intensity per cell was obtained. Neurite length was quantified using the NeuroAnatomy plugin for ImageJ. For mitochondrial analysis, neurons labeled with TOM20 antibody were imaged using a Nikon Ti / CSU-W1 Spinning Disc Confocal microscope with a 100x oil-immersion objective. Mitochondrial morphology was analyzed using the Mitochondria Analyzer plugin for ImageJ.

### Statistics

Statistical analyses were performed using GraphPad Prism 6 (Graph Pad Software Inc., CA). Data were first tested for outliers using the ROUT Method (Q=1%) and identified outliers were excluded from subsequent analyses. Normality of data distribution was assessed using the Anderson-Darling test. Normally distributed data were analyzed with parametric tests, while non-normally distributed data were compared with non-parametric tests. Parametric tests used in this study included the two-tailed unpaired t-test, one-way ANOVA followed by Tukey’s multiple comparison test, and two-way ANOVA followed by Tukey’s multiple comparisons test. Non-parametric tests included the two-tailed Mann–Whitney test and the Kruskal-Wallis test followed by Dunn’s multiple comparisons test. Statistical significance was set at p<0.05 and reported as follows: * for p<0.05, ** for p<0.01, *** for p<0.001, and **** for p<0.0001. ns indicates not significant.

## Acknowledgements

We thank our lab members for their daily help and valuable discussions. This research was supported by grants from Parkinson Schweiz, the Swiss National Science Foundation (1000329),, European Research Council (ERC-StG-311194), Novartis Foundation for medical-biomedical research (19A025), and InnoSuisse (123.355 IP-LS). E.V. was partly supported by the Institute of Genetics and Genomics of Geneva (iGE3). We are also grateful to the team of the Bertarelli Platform for Gene Therapy at EPFL for generating the viral vectors used in this study and advising on their application.

## Author contributions

E.V. and E.N. conceptualized the study. E.V., L.D., and B.S. designed experiments and prepared materials. E.V., L.D. and E.K. performed experiments. E.V., L.D., E.K., O.C. and E.N. analyzed data. E.V., L.D., and E.N., produced visualizations of the results and made figures. E.V, L.D. and E.N. wrote the manuscript.

